# Resting-State fMRI reveals Longitudinal Alterations in Brain Network Connectivity in a Mouse Model of Huntington’s Disease

**DOI:** 10.1101/2022.11.19.517166

**Authors:** Tamara Vasilkovska, Mohit H. Adhikari, Johan Van Audekerke, Dorian Pustina, Roger Cachope, Haiying Tang, Longbin Liu, Ignacio Munoz-Sanjuan, Annemie Van der Linden, Marleen Verhoye

## Abstract

Huntington’s disease is an autosomal, dominantly inherited neurodegenerative disease caused by an expansion of the CAG repeats in exon 1 of the huntingtin gene. Neuronal degeneration and dysfunction that precedes regional atrophy result in the impairment of striatal and cortical circuits that affect the brain’s large-scale network functionality. However, the evolution of these disease-driven, large-scale connectivity alterations is still poorly understood. Here we used resting-state fMRI to investigate functional connectivity changes in a mouse model of Huntington’s disease in several relevant brain networks and how they are affected at different ages that follow a disease-like phenotypic progression. Towards this, we used the heterozygous (HET) form of the zQ175DN Huntington’s disease mouse model that recapitulates aspects of human disease pathology. Seed- and Region-based analyses were performed at different ages, on 3-, 6-, 10-, and 12-month-old HET and age-matched wild-type mice.

Our results demonstrate decreased connectivity starting at 6 months of age, most prominently in regions such as the retrosplenial and cingulate cortices, pertaining to the default mode-like network and auditory and visual cortices, part of the associative cortical network. At 12 months, we observe a shift towards decreased connectivity in regions such as the somatosensory cortices, pertaining to the lateral cortical network, and the caudate putamen, a constituent of the subcortical network. Moreover, we assessed the impact of distinct Huntington’s Disease-like pathology of the zQ175DN HET mice on age-dependent connectivity between different brain regions and networks where we demonstrate that connectivity strength follows a nonlinear, inverted U-shape pattern, a well-known phenomenon of development and normal aging. Conversely, the neuropathologically driven alteration of connectivity, especially in the default mode and associative cortical networks, showed diminished age-dependent evolution of functional connectivity.

These findings reveal that in this Huntington’s disease model, altered connectivity starts with cortical network aberrations which precede striatal connectivity changes, which appear only at a later age. Taken together, these results suggest that the age-dependent cortical network dysfunction seen in rodents could represent a relevant pathological process in Huntington’s disease progression.

## INTRODUCTION

Huntington’s disease is the most prevalent monogenic neurodegenerative disease with an autosomal, dominantly inherited nature.^1,2^ Typically, it develops in young to middle-aged adults and is distinguished by a triad of progressively deteriorating cognitive, psychiatric, and motor signs and symptoms. The origin of Huntington’s disease lies in the abnormal expansion of the CAG repeat above 39 in the huntingtin gene.^3^ This results in a mutated form of the huntingtin protein (mHTT) that contains the expanded polyglutamine (polyQ) stretch. The prevailing hypothesis is that the mHTT protein exhibits toxic gain-of-function properties that lead to dysfunction and subsequent death of neurons.^4^ Vulnerable to this process are basal ganglia structures, especially the striatum and its medium spiny neuron.^4^ However, prominent changes in cortical pathology have been reported, both in anatomy and function, that shed light on the diversity of the Huntington’s disease symptomatology.^5–9^ Before brain atrophy occurs, neuronal properties on the level of both the synapse and circuitry already undergo alterations preceding clinical motor diagnosis.^10–12^ Despite the known genetic background of Huntington’s disease, there is no successful disease-modifying therapy thus far. Towards a better understanding of the disease progression and finding an effective therapeutic strategy, several imaging modalities that hold the potential of biomarkers have been used in both clinical and preclinical research. ^13–15^

One promising imaging modality is resting-state functional MRI (rsfMRI) that measures changes in the blood oxygen level-dependent (BOLD) signal, which indirectly reflects neuronal activity while the brain is at rest. Spontaneous fluctuations in the low frequency (0.01 – 0.1Hz) BOLD signal contain information about the connectivity between distinct brain regions and reveal the brain’s functional architecture, organized in resting-state networks (RSN).^16,17^ Network alterations have already been shown to intersect with neuropathological findings and also to precede and follow clinical manifestation in other neurodegenerative diseases, such as Alzheimer’s and Parkinson’s diseases.^18,19^ In people with Huntington’s disease (PwHD), cross-sectional rsfMRI studies have revealed heterogeneous alterations in different relevant RSNs before and after clinical motor diagnosis.^20^ Most prominent changes are consistently present in the sensory-motor network in PwHD, the visual, default mode, subcortical, and executive networks are mainly affected after clinical diagnosis, however, some studies also report no significant changes in either pre- or different stages post-clinical motor diagnosis.^11,21,22^ Although these clinical findings help describe the course of Huntington’s disease and have the potential to be used in future therapeutic strategies, conflicting evidence suggests the need for a longitudinal study to understand the evolution of functional connectivity (FC) alterations of these networks consequential to the development and progression of Huntington’s disease.

Studies in rodent models of Huntington’s disease have affirmed the relevance of both the cortical and striatal networks where neuronal activity impairments are present in both the cortical projection and medium spiny neurons across ages.^23–27^ Cortical circuitry alterations in Huntington’s disease have been gaining more attention where electrophysiological findings have shown that, under sensory stimulation, cortical areas such as the motor and sensory cortex demonstrate aberrant activity in early manifest states.^25,26^ However, whole-brain functional connectivity changes in rodent models have not been well investigated, hence the lack of assessment of the macroscopic network alterations along the Huntington’s disease-like phenotype progression.

Here we aimed to assess the changes in resting-state network FC using the knock-in zQ175 delta-neo (DN) heterozygous (HET) mouse model of Huntington’s Disease, which exhibits cellular, behavioral and cognitive abnormalities, where motor alterations follow temporal progression from 6 months onwards. ^28–30^ We acquired rsfMRI and assessed FC at four different ages: 3, 6, 10 and 12 months, in the zQ175DN HET, and age-matched wild-type (WT) mice. We hypothesized that connectivity changes in the large-scale brain networks, especially involving the striatum, are altered at an early age in the zQ175DN HET mice. Moreover, we hypothesized that the already established mHTT progressive accumulation^30^ leads to continuous worsening in the cortico-cortical and cortico-striatal connectivity of the zQ175DN HET mice.^30^

## MATERIALS & METHODS

### Animals

Two cohorts of male, age-matched, WT and HET (cohort 1: WT (*n* = 18), HET (*n* = 19); cohort 2 WT (*n* = 16), HET (*n* = 19)) zQ175DN KI littermates (C57BL/6J background, CHDI-81003019, JAX stock #029928) were obtained from the Jackson Laboratory (Bar Harbor, ME, USA). Animals were single-housed in individually ventilated cages with food and water *ad libitum* and continuous monitoring for temperature and humidity under a 12h light/dark cycle.

The animals were kept in the animal facility for at least a week to acclimatize to the current conditions before the experimental procedures. All experiments and handling were done in accordance with the EU legislation regulations (EU directive 2010/63/EU) and were approved by the Ethical Committee for Animal Testing UAntwerp (ECD # 2017-09).

The zQ175DN KI (without a floxed neomycin cassette) mouse model has the human HTT exon 1 substitute for the mouse *Htt* exon 1 with ~180-220 CAG repeats long tract. This model is a modified version of the zQ175 KI^28^ where the congenic C57BL6J is used as the strain background.^31^ The first motor deficits are observed at 6 months, marked as an onset of phenoconversion.^29^ The HET form of zQ175DN has a slow progression reflected in the increase of mHTT aggregation from 3 until 12 months, initially appearing in the striatum at 3 and later on in the cortex at 8 months of age.^30^ A longitudinal rsfMRI study was performed in the first cohort, in both the zQ175DN HET and their age-matched WTs^32^, at different ages following Huntington’s disease-like phenotypic progression: 3, 6 and 10 months. rsfMRI data in the second cohort were obtained at 12 months of age.

### rsfMRI acquisition

MRI scans were acquired on a 9.4 T Biospec system with a four-element receive-only mouse head cryoprobe coil (Bruker Biospin MRI, Ettlingen, Germany) and a volume resonator for transmission. Prior to the whole-brain rsfMRI scan, a B0 field map was acquired to measure the magnetic field inhomogeneities after which local shimming was performed within an ellipsoid volume, covering the middle portion of the brain. The dynamic BOLD resting-state signals were measured with a T2*-weighted single-shot Echo-Planar Imaging (EPI) sequence (field of view (FOV) (27×21) mm^2^, matrix dimensions (MD) [90×70], 12 horizontal slices of 0.4mm, voxel dimensions (300×300×400) μm^3^, flip angle 60°, TR 500ms, TE 15ms, 1200 repetitions). After the rsfMRI scan was acquired, a 3D anatomical scan was obtained using the 3D Rapid Acquisition with Refocused Echoes (RARE) sequence with FOV (20×20×10) mm^3^, MD [256×256], 128 horizontal slices of 0.3mm, (78×78×78) μm^3^, TR 1800ms, TE 42ms, pixel dimensions.

During rsfMRI scans, mice were initially anesthetized with 2% isoflurane (Isoflo^®^, Abbot Laboratories Ltd., USA) in a mixture of 200ml/min O_2_ and 400ml/min N_2_. After the animal was stabilized, a subcutaneous bolus injection of medetomidine (0.075 mg/kg; Domitor, Pfizer, Karlsruhe, Germany) was applied followed by a gradual decrease to 0.5% isoflurane over the course of 30 minutes which was kept at this level throughout the whole experiment.

Meanwhile, a continuous infusion of medetomidine (0.15 mg/kg/h), starting 10 minus post bolus medetomidine injection, was applied in combination with the isoflurane, an established protocol for rsfMRI in rodents.^33,34^ Throughout the duration of the experiment, all physiological parameters (breathing rate, heart rate, O2 saturation, and body temperature) were kept under normal conditions.

### rsfMRI image preprocessing

rsfMRI repetitions for each session were realigned to the first image with a 6-parameter rigid-body spatial transformation estimated with the least-squares approach. Next, a study-based template was generated. To create an unbiased template, in cohort 1, we used the individual 3D RARE images from 1/2 of each group from each time point. In cohort 2, all 3D RARE images from both groups were used to generate the study template. The rsfMRI data were co-registered to their respective subject 3D RARE image. The 3D RARE images were normalized to the common study template. Spatial transformation parameters were also estimated between the study-based template and an in-house C57BL/6 mouse brain atlas. All the rsfMRI data were spatially normalized to the in-house C57BL/6 atlas, combining all estimated transformation parameters: (1) rsfMRI to 3D RARE, (2) 3D RARE to common study template, and (3) common study template to in-house C57BL/6 atlas. In-plane smoothing was applied using a Gaussian kernel with full width at half maximum of twice the voxel size followed by motion vector regression based on the parameters generated during realignment. These steps were performed using Statistical Parametric Mapping (SPM) using SPM12 (Wellcome Centre for Human Neuroimaging, London, UK). Using a whole-brain mask, images were further filtered (0.01-0.2Hz) with a Butterworth band-pass filter where five repetitions from both the beginning and the end of the image series were removed before and after filtering to eliminate transient effects. Finally, quadratic detrending (QDT), voxel-wise global signal regression (GSR), and normalization to unit variance were applied. All analysis steps were performed with MATLAB R2021a (Mathworks, Natick, MA) and template creation was done using Advanced Normalization Tools (ANTs).

### Functional connectivity (FC) analysis

To assess FC, we performed connectivity analysis on three different levels: Region of Interest (ROI)-, network- and seed-based FC. ROI-based FC was performed on selected ROIs that pertain to four different networks: the Default Mode-like Network (DMLN), associative cortical network (ACN), Lateral Cortical Network (LCN), and the Subcortical Network (SuCN). The network-based FC was carried out by pooling the FCs of ROI pairs that belong to the specific networks (within-network FC) or a pair of networks (between-network FC). Moreover, to investigate the brain-wide functional topography, we further performed seed-based FC analysis, where we used several representative regions from each network to evaluate the connectivity of each seed with the rest of the brain.

### ROI- and network-based FC

We selected 26 unilateral ROIs (both left (L) and right (R) hemispheres for each region) from an in-house C57BL6 mouse atlas^35^ that represent the main hubs of several relevant large-scale networks.^36,37^ The abbreviations for each region are presented in Table 1. For each subject and each region, we extracted the time series of the region-averaged BOLD signal. Pearson correlation coefficients were calculated between the BOLD signal time series of each pair of regions. These correlations were Fisher z-transformed, thus obtaining subject-wise FC matrices. FC within each brain network was calculated by averaging the pairwise correlations between the regions of that network. Similarly, between-network FC was calculated by averaging correlations between the regions of each network. All the above scores were computed for each subject individually at each age.

**Table 1.** Brain regions and Network abbreviations.

| Network | Region |
| --- | --- |
| Default Mode-Like Network (DMLN) | CgCtx Cingulate cortex |
|  | RspCtx Retrosplenial cortex |
|  | PrlCtx Prelimbic cortex |
| Associative Cortical Network (ACN) | VCtx Visual cortex |
|  | AuCtx Auditory cortex |
| Lateral Cortical Network (LCN) | S1Ctx Somatosensory cortex 1 |
|  | S2Ctx Somatosensory cortex 2 |
|  | MCtx Motor cortex |
|  | InsCtx Insular Cortex |
|  | FrACtx Frontal Association Cortex |
| Subcortical Network (SuCN) | mCPu Caudate Putamen, medial division |
|  | ICPu Caudate Putamen, lateral division |
|  | PirCtx Piriform cortex |

### Seed-based FC

Seeds (4×4×1 voxels) were drawn at the center of representative regions that pertain to the four large-scale networks. Seeds were drawn on the left hemisphere regions using the atlas parcellation to guide their placement. The selected regions were representative of each of the large-scale networks (Table 1). The BOLD signal time series were extracted for each seed and used in the multiple regression model of the BOLD signal time series of every voxel, thus producing seed-specific individual FC maps.

### Statistical analysis

For ROI- and network-based FC, significant connections within each group (One-sample T-test, *p* ≤ 0.05, False Discovery Rate (FDR) corrected) were identified, and between-group FC differences were tested for FC pairs found to be significantly different in at least one of the groups (Two-sample T-test, FDR corrected, *p* ≤ 0.05). ROI- and network-based FC statistics were performed in MATLAB R2021a. For Seed-based FC, group-level seed-based FC maps were generated for both the WT and HET group (One-sample T-test, FDR corrected, p ≤ 0.05, cluster size (k) ≥ 10). A union of the group-level masks of voxels which are statistically significantly correlated with the seed region was applied when calculating between-group differences for each age (Two-sample T-test, FDR corrected, p≤0.05, k≥10) so that only voxels correlating with the seed region in WT and/or in HET were analyzed. Voxel-level statistics for the seed-based analysis was performed using SPM12. For visual representation, the T-statistics were upsampled and transferred onto the high-resolution Australian mouse brain MRI anatomical image^38^. These visualizations were produced with MRIcroGL (McCausland Center for Brain Imaging, University of South Carolina, USA).

In cohort 1, a longitudinal assessment of changes in FC for selected ROI pairs as well as within and between networks was performed using a Linear Mixed Model (LMM) in JMP^®^ (Version 16, SAS Institute Inc., Cary, NC, 1989 – 2021). We set age, genotype, and age*genotype interaction as fixed effects in the LMM and added a random slope model with the subject as a random intercept and age as a random slope. In the case when no significant age*genotype interaction was present, the interaction was removed from the LMM and the model was recalculated using genotype and age and those p values are reported. A posthoc test was further applied using Tukey honestly significant difference (HSD) with *p* ≤ 0.05. Graphical representation of the data was obtained using GraphPad Prism (version 9.2.0 for Windows, GraphPad Software, San Diego, California USA, www.graphpad.com). All data are represented with an interquartile range plot and subject values.

## DATA AVAILABILITY

The in-house C57BL6 mouse atlas used in this study is publicly available https://www.uantwerpen.be/en/research-groups/bio-imaging-lab/research/mri-atlases/c57bl6/. The data that support the findings of this study are available from the corresponding author upon reasonable request.

## RESULTS

### Reduced FC within DMLN and ACN regions at 6 but not 3 months of age

At 3 months, in both groups, positive inter-hemispheric connectivity between homologous regions within all four networks was present (Fig. 1A). In addition, a positive correlation was also found between regions of the DMLN and ACN, while a negative correlation of regions pertaining to these two networks with LCN regions was observed. Moreover, the lCPu was positively correlated with S1Ctx and S2Ctx from the LCN while negatively correlated with regions of the DMLN and ACN. Genotype comparison showed no significant differences at this premanifest state (*p*>0.05, FDR corrected, Fig. 1B). At 6 months, in both WT and zQ175DN HET, inter- and intrahemispheric positive FC was present in each network (Fig. 1C). Moreover, the positive FC between regions of the DMLN and ACN and between LCN and SuCN, as well as the anti-correlations of these network region pairs, persisted as observed at 3 months. Between-group comparison revealed significantly decreased connectivity in the zQ175DN HET mice (*p*≤0.05, FDR corrected, Fig. 1D) in several pairs of regions within the DMLN: RspCtx (L - R), RspCtx (L&R) – CgCtx (L), PrlCtx (R) – RspCtx (L&R), in the ACN: VCtx (L – R), VCtx (L) – AuCtx (L) and between DMLN and ACN: AuCtx (L) – CgCtx (L&R), AuCtx (L) – RspCtx (L&R).

**Figure 1.**
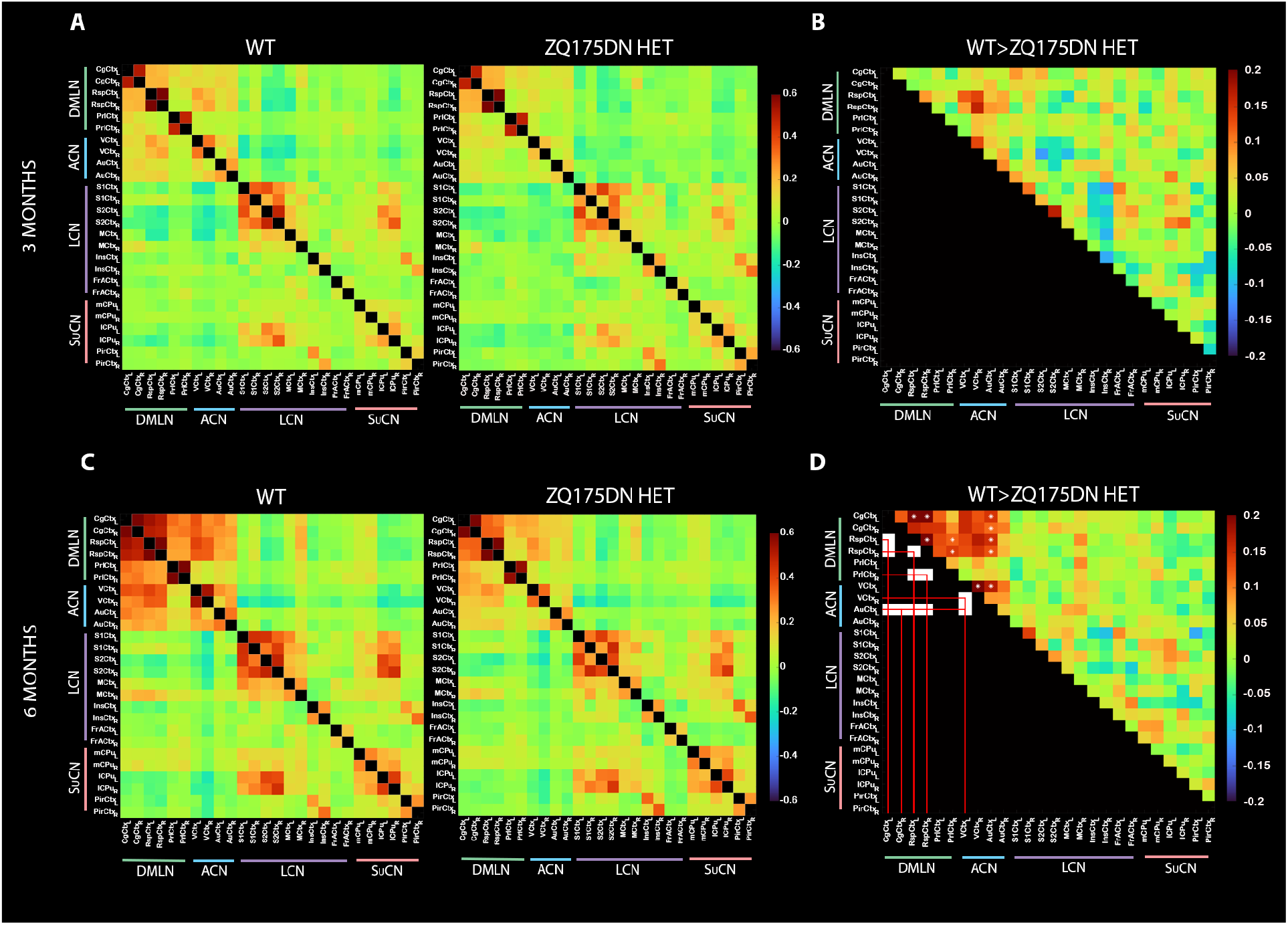
ROI-based FC shows decreased connectivity within and between DMLN and ACN regions in the zQ175DN HET at 6 months of age. **(A, C)** Mean z-transformed correlation (mirrored) matrices of each group at 3 (top row) and 6 months of age (bottom row); red/orange colors represent positively correlated connectivity, green color indicates very low to no connectivity between regions, and dark/light blue colors represent negatively correlated connectivity. **(B, D)** The upper triangle shows the FC differences between groups for 3 and 6 months of age, respectively. Red/orange represents lower positive or higher negative FC between a pair of regions in the zQ175DN HET compared to WT while dark/light blue represents higher positive or lower negative FC in the zQ175DN HET compared to WT. White squares in the lower triangle and asterisk in the upper triangle indicate significant group differences in FC based on a two-sample T-test (*p* ≤ 0.05, FDR corrected) only performed on connections that demonstrated a significant FC in at least one group based on a one-sample T-test.

### Decreased connectivity in LCN and SuCN regions at 10 months of age

At 10 months, in both WT and HET, within all four networks, there was a positive FC within each pair of regions (Fig. 2A). Moreover, a continued positive FC of DMLN regions with ACN regions and with regions from the LCN with the SuCN, was observed as it was in other ages (Fig. 1). Inter-genotype comparisons showed a significant decrease in FC in the zQ175DN HET mice between PrL (L – R) and AuCtx (R) – VCtx (R) (*p*≤0.05, FDR corrected, Fig. 2B). At 12 months, besides the positive FC within and between regions of the DMLN, ACN and SuCN in both groups, in the LCN, there was no interhemispheric FC present in the S1Ctx and S2Ctx (Fig. 2C). This was reflected in a positive intrahemispheric FC of these with all other regions of LCN, but no interhemispheric FC was observed. Interestingly, despite the positive homotopic FC for mCPu and lCPu of the SuCN, both regions showed only positive intrahemispheric correlations with the other regions of SuCN. Notably, the zQ175DN HET showed a significantly decreased interhemispheric FC between RspCtx (L – R), AuCtx (L – R), S2Ctx (L – R), InsCtx (L) – S2Ctx (R), lCPu (L) – InsCtx (R), lCPu (R) – InsCtx (L) and intrahemispheric lCPu (R) – PirCtx (R) (*p*≤0.05, FDR corrected, Fig. 2D). Besides the overall decrease in the positively correlated pairs, at this age, there was also a significant decrease in the zQ175DN HET in the negative FC between MCtx (L) – VCtx (R).

**Figure 2.**
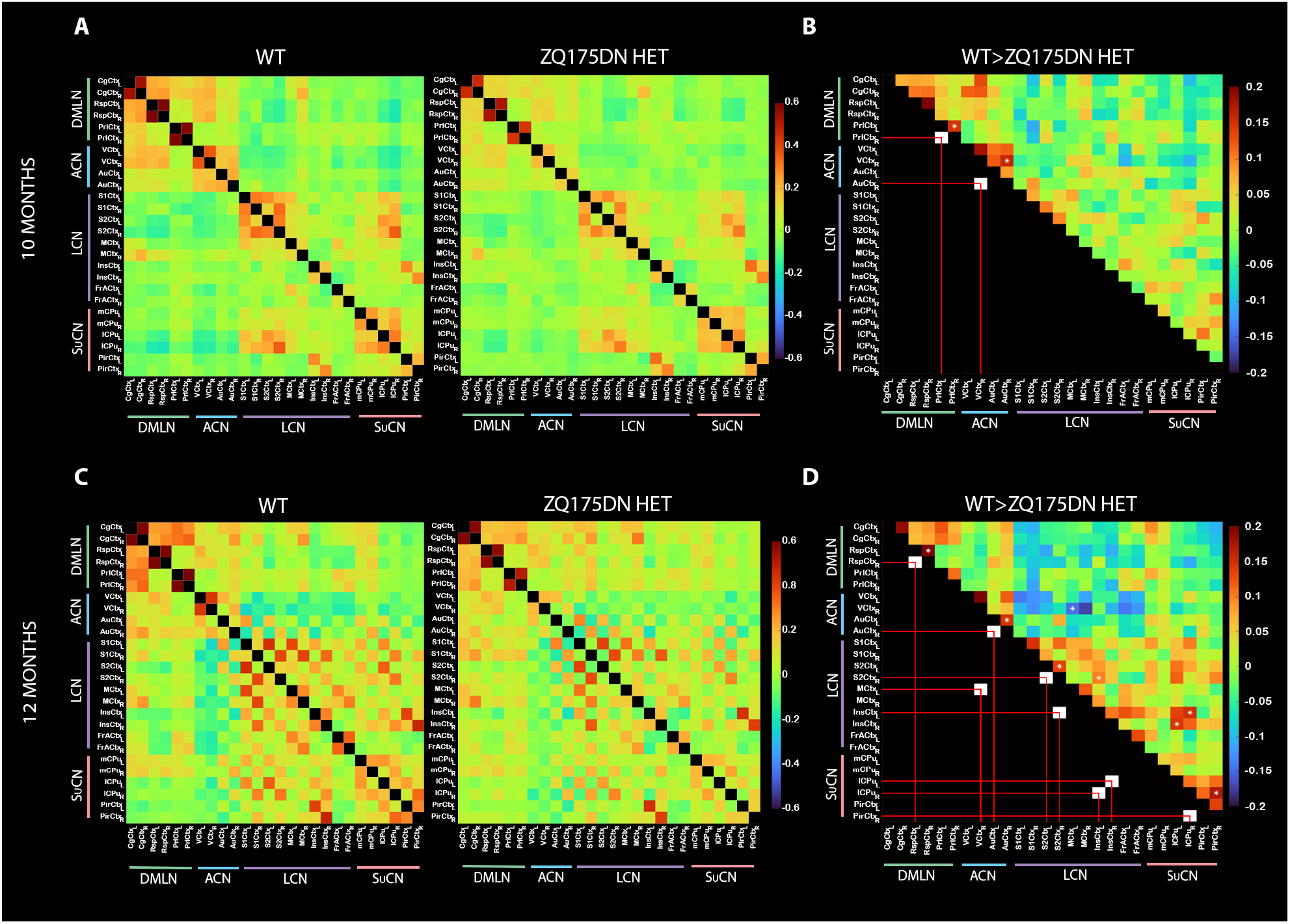
Reduced FC in brain networks at 10 and 12 months of age in the zQ175DN HET mice. **(A, C)** Mean z-transformed correlation (mirrored) matrices of each group at 10 (top row) and 12 months of age (bottom row); red/orange colors represent positively correlated connectivity, green color indicates very low to no connectivity between regions and dark/light blue colors represent negatively correlated connectivity **(B, D)** Upper triangle shows the between-group FC differences for 10 and 12 months of age, respectively. Red/orange represents lower positive or higher negative FC between a pair of regions in the zQ175DN HET compared to WT while dark/light blue represents higher positive or lower negative FC in the zQ175DN HET compared to WT. White squares in the lower triangle and asterisk in the upper triangle indicate significant group differences in FC based on a two-sample T-test (*p* ≤ 0.05, FDR corrected) only performed on connections that demonstrated a significant FC in at least one group based on a one-sample T-test.

### Decrease in cortical and hippocampal connectivity at 6 months of age

Seed-based connectivity maps for each selected seed were obtained for both groups at each age, however, we found significant inter-genotype difference only at 6 months (Fig. 3), whereas for 3, 10, and 12 months we found no significant FC difference for any of the seeds (data not shown).

**Figure 3.**
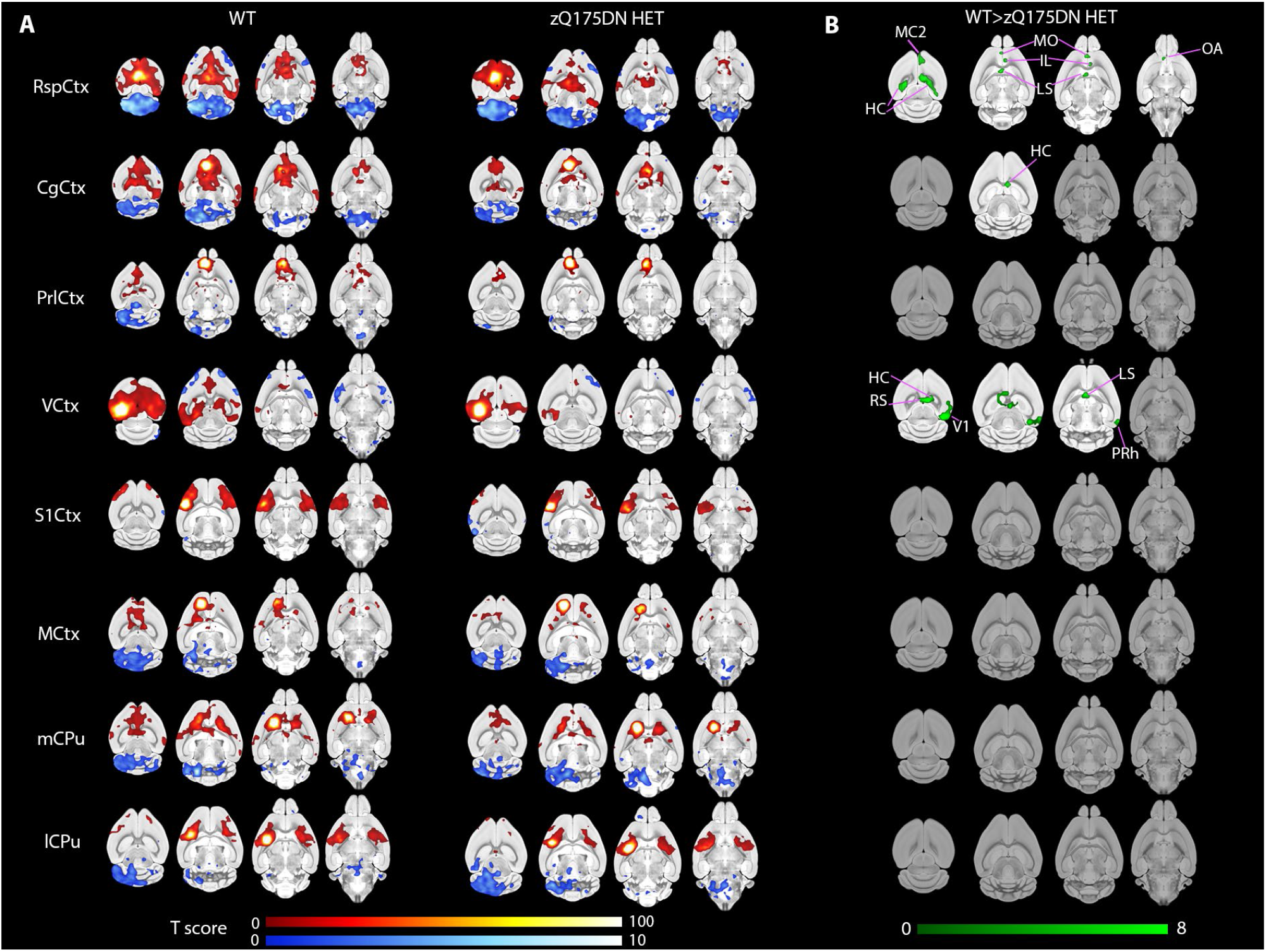
Seed-based FC analysis reveals reduced connectivity in the zQ175DN HET at 6 months of age. **(A)** Group-level T-statistical maps of significantly positively (red) or negatively (blue) correlated regions with the unilateral seeds in both WT and HET **(B)** Voxel-wise significant differences in connectivity WT>HET for each seed; T-statistics are FDR corrected for *p* ≤ 0.05 and cluster size (k ≥ 10) corrected for *p* ≤ 0.05; color scales represent T-values; MC2 - Motor Cortex 2, HC – Hippocampus, MO – Medial Orbital Cortex, IL – Infralimbic Cortex, LS – Lateral septal nucleus, OA – Olfactory Area, PRh – Perirhinal Cortex.

At 6 months, using seed-based analysis, we obtained voxel-wise connectivity maps in both groups for several seeds that are pertaining to DMLN, ACN, LCN, and SuCN. As part of the DMLN, both the RspCtx and the CgCtx seeds showed positive FC with other DMLN constituents as well as with the dentate gyrus (DG), medial orbital cortex (MO), Infralimbic Cortex (IL), Lateral septal nucleus (LS), and Olfactory Area (OA), while a negative FC with the cerebellum (CB) and medulla (MY) was present (Fig. 3A). Inter-group comparison showed, in the zQ175DN HET mice, decreased FC of the RspCtx with the positively correlated regions, while the decreased FC of the CgCtx seed was only with the HC (Fig. 3B). The VCtx seed, part of the ACN, was positively correlated with the contralateral VCtx and with RspCtx, CgCtx, MCtx, HC, and LS while negatively correlated with some portions of S1Ctx (Fig. 3A). A significantly decreased FC in the zQ175DN HET of VCtx with HC, RspCtx, contralateral VCtx, LS, and the perirhinal cortex was observed (Fig. 3B). However, seeds in S1Ctx and MCtx as part of LCN and the CPu from SuCN (Fig. 3A) showed no significant difference in FC between genotypes (Fig. 3B).

### Network connectivity decreases from 6 to 12 months of age

In addition to the inter-regional FC differences observed in zQ175DN HET at different states, we aimed to also assess the disease-related alterations on a network level. As in the case of inter-regional FC, there was no significant difference in network FC between genotypes at 3-months of age (Supplementary Fig. S1B, top row). At 6 months, in both groups, the positive within DMLN, within ACN and DMLN – ACN, and LCN - SuCN connectivity was present (Fig. 4A, top row). Genotypic differences at this age include decreased FC within DMLN, within ACN, and between DMLN – ACN in the zQ175DN HET, in line with the observed ROI-level FC differences (Fig. 1B). At 10 months, only a decrease within the ACN was found (Supplementary Fig. S1B, bottom row). At 12 months, the within-network, DMLN – ACN, and LCN – SuCN FC continued to be positive as in earlier ages in both groups. However, the LCN-DMLN anti-correlation, observed in both groups at earlier ages as well as in WT at 12 months, was no longer present in the zQ175DN HET group as a positive LCN-DMLN FC was observed (Fig. 4A, bottom row). Between-group comparisons revealed a significantly decreased within ACN and within SuCN FC and a decreased negative FC between LCN and ACN in the zQ175DN HET (Fig. 4B, bottom row).

**Figure 4.**
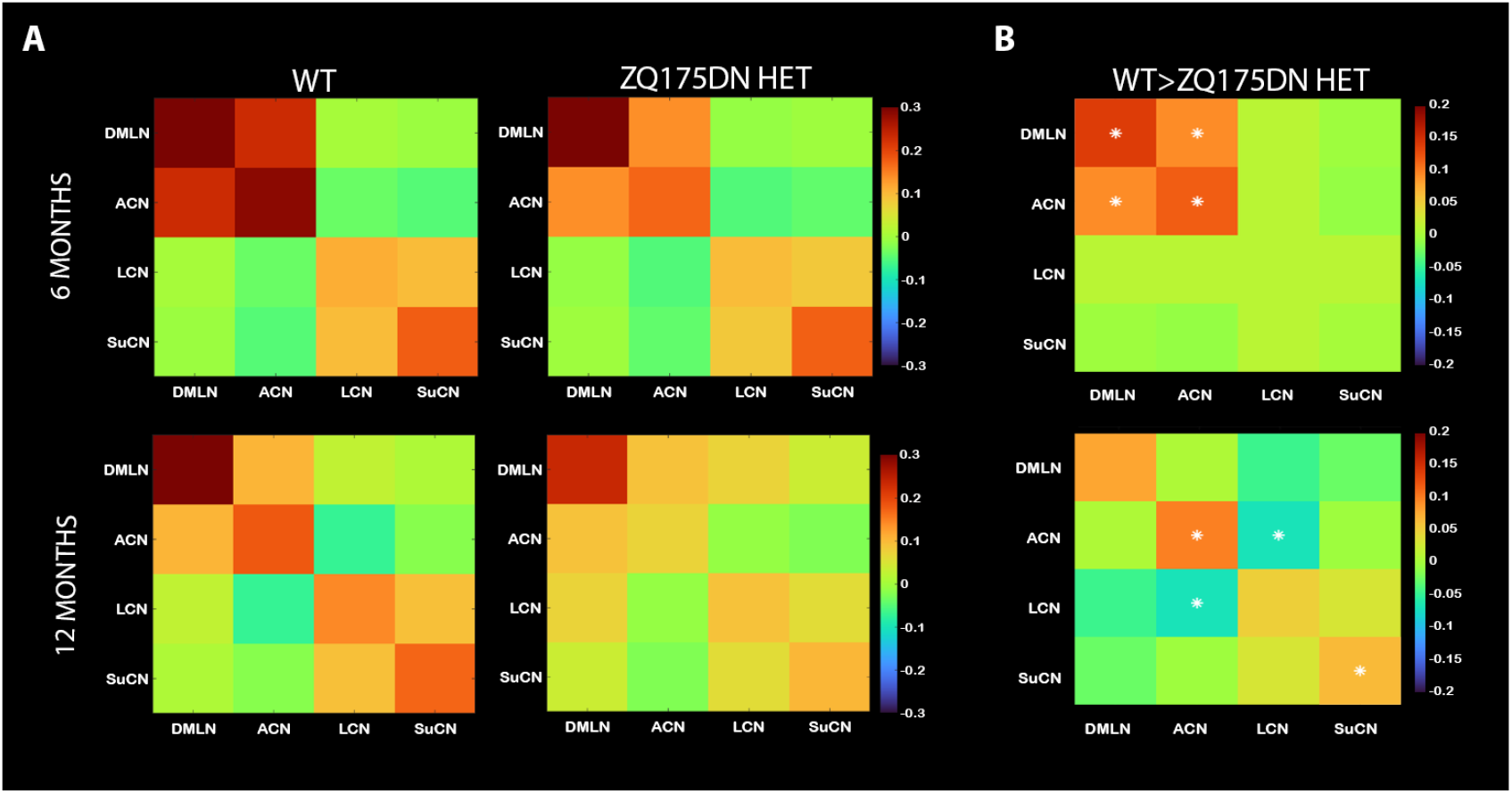
Network FC decreases at 6 and 12 months of age in zQ175DN HET and WT mice. **(A)** Mean z-transformed correlation (mirrored) matrices of each group at 6 (top row) and 12 months of age (bottom row) – within network FC is shown along the diagonal of the matrix; red/orange colors represent positively correlated connectivity, green color indicates very low to no connectivity between regions and dark/light blue colors represent negatively correlated connectivity. **(B)** Between-group differences for 6 and 12 months of age. Red/orange represents lower positive or higher negative FC between a pair of regions in zQ175DN HET compared to WT while dark/light blue represents higher positive FC or lower negative FC in zQ175DN HET compared to WT. Asterisk indicates significant group differences in FC based on a two-sample T-test (*p* ≤ 0.05, FDR corrected) only performed on connections that demonstrated a significant FC in at least one group based on a one-sample T-test.

### Age-dependent, region-based FC changes along phenotypic progression

In the cross-sectional comparisons, we observed differences in connectivity strength in different ages in both groups. Additionally, as development and normal aging have been shown to have an impact on FC^39^, we sought to investigate the disease effect on age-dependent connectivity changes of several different FC ROI pairs in this model.

Overall, in the WT group, in the majority of the region pairs, there was an increase in FC from 3 to 6 months of age, followed by a decreased FC from 6 to 10 months of age (Fig. 5). This non-linear inverted U-shape has been reported as a normal product of aging. To understand if this is the case in the zQ175DN HET as well, we first assessed the interhemispheric FC of four representative regions from each network (Fig. 5, top row). In all four regions, there was no significant interaction between age and genotype (Table 2). However, there was a significant age effect following the same inverted U-shape as in the WT group, with the exception of the S2Ctx which showed no significant change in FC from 3 to 6 months (Fig. 5, top row). Interestingly, a significant genotype effect was present in RspCtx (*p* = 0.0001), S2Ctx (*p* = 0.0171) and VCtx (*p* = 5.0E-06) but not in the lCPu (*p* = 0.7144). Thus, while the inverted U-shape age effect was present in the inter-hemispheric FC of three out of four network representatives in both genotypes, there was an overall, age-non-specific reduction in FC in the zQ175DN HET group in these ROIs. In addition, we assessed the FC evolution between regions from different networks, in an ROI pair of the DMLN - CgCtx with RspCtx, of the ACN – AuCtx with VCtx, and between DMLN and ACN - AuCtx with RspCtx (Fig. 5, bottom row). A significant interaction of age and genotype was only present in the Cg – RspCtx connection (*p* = 0.0461), where the posthoc comparisons revealed a decreased FC at 6 months in zQ175DN HET (*p* = 0.0118). Furthermore, as opposed to the inverted U-shape FC in the WT, with a significant increase in FC from 3 to 6 months (*p* = 3.00E-05) followed by a decrease from 6 to 10 months (*p* = 4.60E), zQ175DN HET showed a decrease of FC only from 6 to 10 months (*p* = 0.016). In the Au – VCtx and Au – RspCtx pairs, there was a significant genotype effect (see Table 2) but only Au – RspCtx showed an age effect in the form of an inverted U-shape (Fig. 5, bottom row). Additionally, as the somatosensory regions have been shown as one of the critically affected areas in Huntington’s disease^8^, we explored the disease effect on age-dependent changes within an ROI pair of the LCN – between S2Ctx and MCtx where no interaction nor a significant age or genotype effect was found (Fig. 5, bottom row).

**Figure 5.**
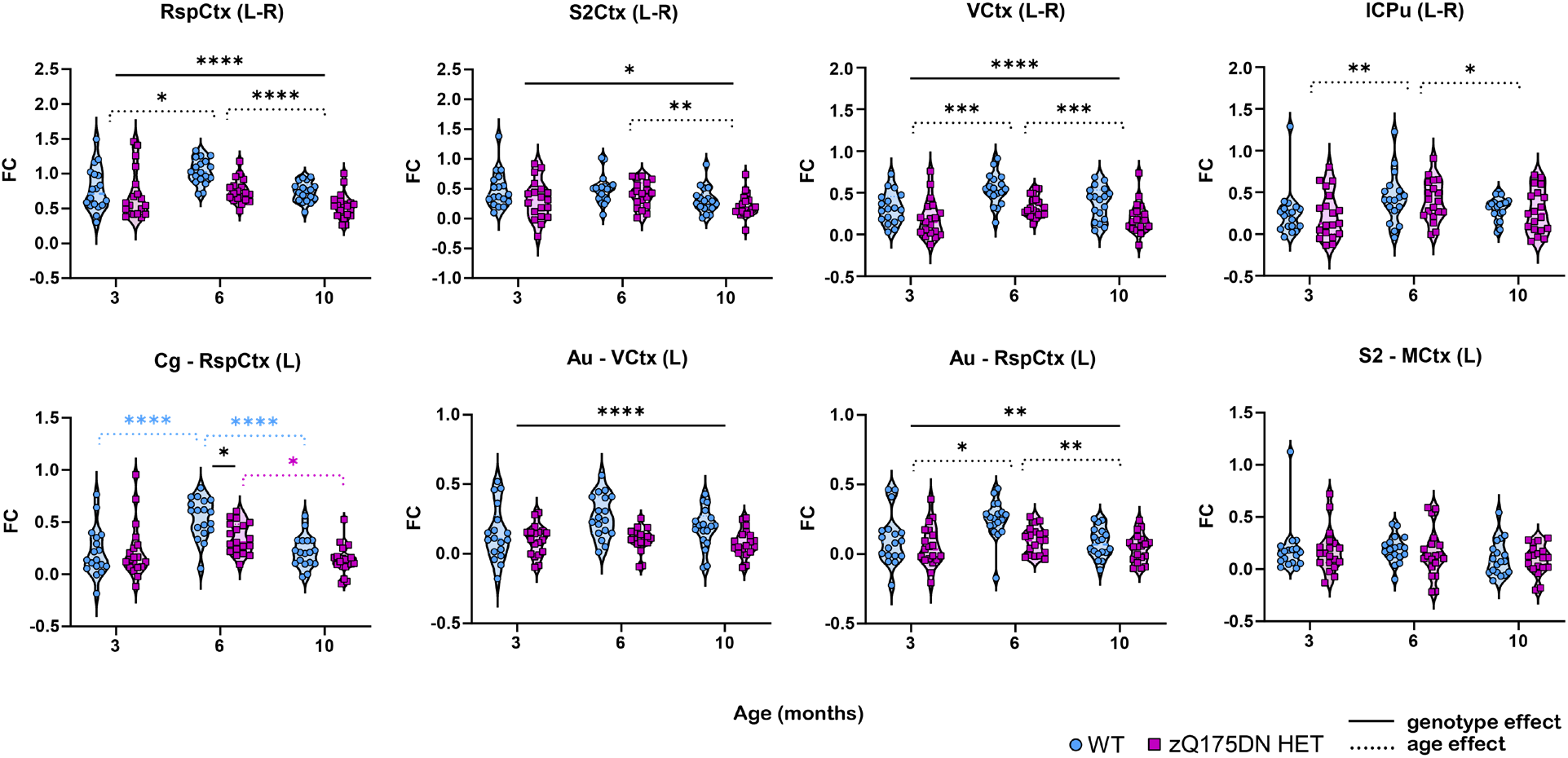
Age-dependent change in pairwise FC of a subset of regions in WT and zQ175DN HET mice. Plots demonstrate the connectivity change over time for different pairs of regions: RspCtx (L-R), S2Ctx (L-R), VCtx (L-R), lCPu (L-R), Cg – RspCtx (L), Au - VCtx (L), Au – RspCtx (L), S2 - MCtx (L) for both genotypes. Data are presented as an interquartile range of distribution with subject values in both WT (blue circle) and HET (magenta square). Full lines represent the main genotype effect and genotype differences at a certain time point; Dashed lines represent the overall age effect (black) or within WT (blue) and within zQ175DN HET (magenta). Significant difference after Tukey HSD * *p* ≤ 0.05, ** *p* ≤ 0.01, *** *p* ≤ 0.001, **** *p* ≤ 0.0001

### Age-susceptible network alterations across different phenotypic states

As genotype effects were observed on age-dependent FC changes in several ROI pairs, we also investigated whether the disease impacts the normal aging of network-level FC. Initially, we assessed the intra-network FC for all four networks (Fig. 6, top row). No significant interaction between age and genotype was found in any of the four networks, however, a significant age effect was present (see Table 3), with the already observed non-linear trend (Fig.6, bottom row). A significant genotype effect was found only in the DMLN and ACN. Subsequently, we evaluated the impact of the disease on the inter-network connectivity, especially in the networks that were shown to be impacted in the zQ175DN HET group (Fig. 4B, Supplementary Fig.S1B) and are either positively (DMLN with ACN and LCN with SuCN) or negatively (LCN with DMLN and LCN with ACN) correlated (Fig. 6, bottom row). No interaction of age and genotype was present between any of the networks (see Table 2), but there was a significant age effect in the DMLN – ACN and LCN – SuCN pairs, following the same non-linear trend, and a significant genotype effect only found in the DMLN – ACN FC (Fig. 6, bottom row).

**Figure 6.**
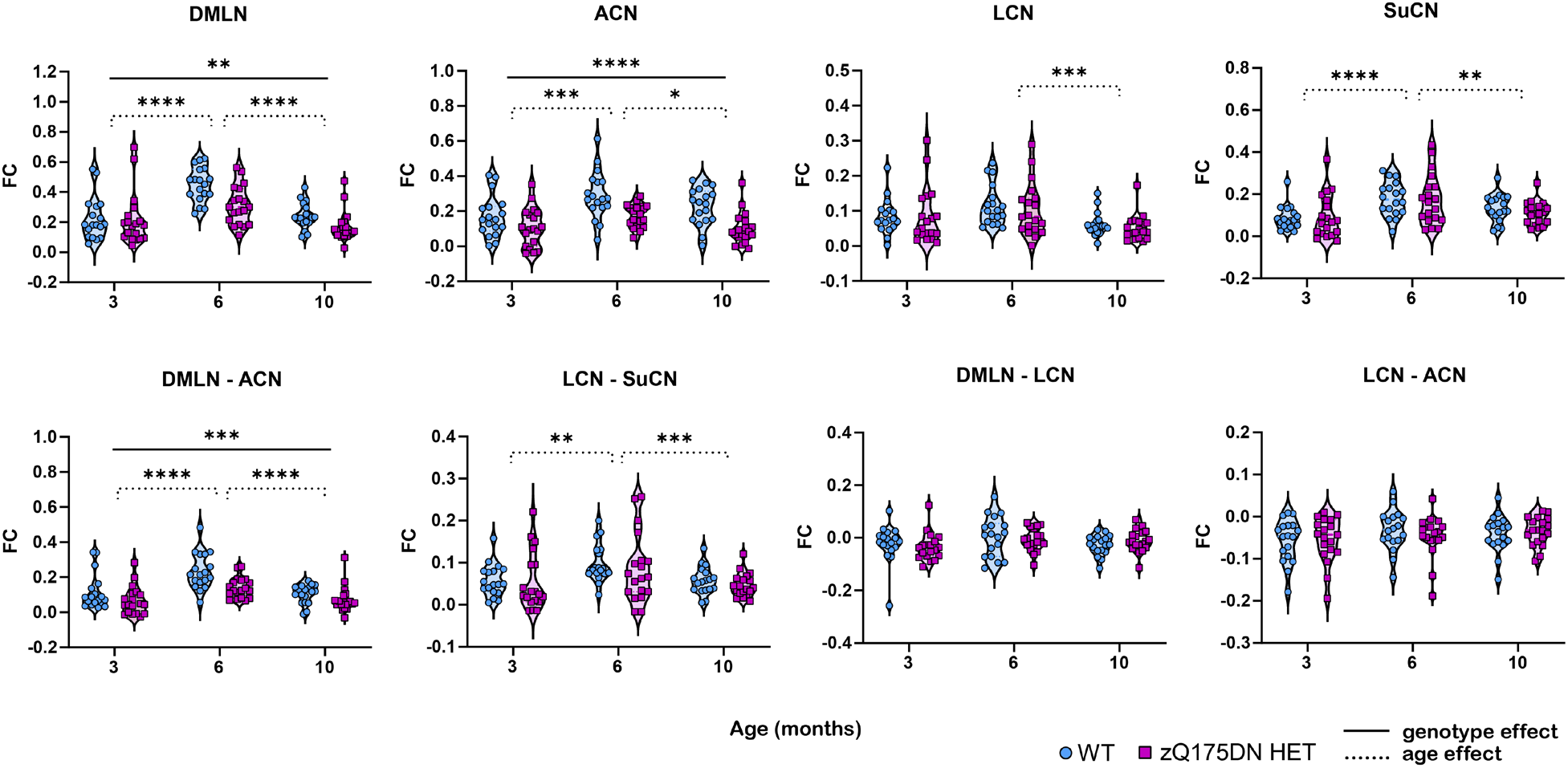
Age-dependent change in network FC in both WT and zQ175DN HET mice. Plots demonstrate the connectivity change over time for different networks: intra-DMLN, ACN, LCN, SuCN, and between DMLN – ACN, LCN – SuCN, DMLN – LCN, LCN – ACN for both genotypes. Data are presented as an interquartile range of distribution with subject values in both WT (blue circle) and zQ175DN HET (magenta square). Full lines represent the main genotype effect and genotype differences at a certain time point; Dashed lines represent the overall time effect (black) or within WT (blue) and zQ175DN HET (magenta). Significant difference after Tukey HSD * *p* ≤ 0.05, ** *p* ≤ 0.01, *** *p* ≤ 0.001, **** *p* ≤ 0.0001

## DISCUSSION

To our knowledge this study is the first to investigate longitudinal changes in large-scale brain network functional connectivity across disease progression in a Huntington’s disease animal model. At 3 months, our findings showed no significant difference in either region or network FC in the zQ175DN HET compared to WT. At both 6 and 10 months, connectivity reductions between regions of the default mode-like and associated cortical network were continuously present in the zQ175DN HET mice. Interestingly, at 12 months of age, a shift towards decreased connectivity in the lateral and subcortical regions appeared, where the lateral CPu, the most vulnerable region in Huntington’s disease, exhibited reduced cortical connectivity. Moreover, our results demonstrated age-dependent connectivity changes as a byproduct of normal aging in the zQ175DN HET mice, in the form of an inverse U-shape, as observed in the WT group. The neuropathologically driven impact on the age-dependent FC evolution was present in the default mode and associative cortical networks with diminished connectivity in the zQ175DN HET mice. In addition, at 6 months, reduced cingulate – retrosplenial cortex connectivity interfered with the normal FC progression in the zQ175DN HET mice.

The retrosplenial, cingulate, pre- and infralimbic, orbital cortex, and hippocampus are the constituents of the DMLN, a rodent analog to the human default mode network (DMN), which represents correlated activity between these brain regions in the absence of a goal-oriented task.^36,40–42^ Many studies have accentuated the relevance of DMN, especially in neurological disorders. ^43–46^ Our results from both seed- and ROI-based FC analyses have demonstrated consistently a reduced FC within DMLN connectivity in the zQ175DN HET mice, including the retrosplenial and cingulate cortices, and the medial prefrontal cortex (mPFC) regions such as orbital, pre- and infralimbic cortices^47^, present in an early disease-like phenotypic state. These findings are consistent with widespread DMN alterations observed in PwHD.^21,48,49^

At 6 months, with a seed in the retrosplenial cortex, we observed decreased FC with the motor cortex 2, hippocampus, lateral septum, and the mPFC regions. Interestingly, the relationship in mice between the retrosplenial and motor cortex is known to be involved in sensorimotor processing and motor control.^50^ Hence, impaired FC between those regions might potentially explain the motor abnormalities that start at 6 months of age. The retrosplenial cortex and the lateral septal nucleus are regions that are important in memory processing and hippocampal efferent projections to these areas are critical in cognitive processes.^51^

Noteworthy is that at 6 months, cognitive deficits are present in this Huntington’s Disease mouse model and others,^52,53^ which is in line with the well-known cognitive and memory deficits described in PwHD.^54–60^

Alterations in the auditory system have been reported in both preclinical and PwHD.^61–63^ In line with this, our findings showed, at 6 months, a decreased FC between the auditory with the cingulate, retrosplenial, and visual cortices in the zQ175DN HET mice. The cingulate is part of the auditory cortical network while the retrosplenial cortex is relevant in auditory memory-related processes.^64,65^ Most importantly, the decreased FC in the associative network, comprised of the auditory and visual cortices, indicates impaired visual and auditory processing at this age; this supports recent findings of altered dynamics of the sensory and motor cortices of the zQ175 HET model when using auditory and visual sensory stimulation at the same age of 6 months.^26^ In addition, the reduced FC of the visual cortex with memory-related regions (HC, LS, PRh) and the retrosplenial cortex, accentuates our findings of marked cognitive and sensory cortical connectivity alterations at this early Huntington’s disease-like phenotypic state.

Our findings show a shift in network alteration towards the lateral cortical and subcortical network regions in the zQ175DN HET mice at 12 months of age. Compared to the early changes where we observed altered FC in multiple connections from both DMLN and the associative network, at this age, only interhemispheric connectivity of retrosplenial and auditory cortices showed reduced FC, from each network respectively. The most interesting finding by this age, are the reduced changes related to these networks and the more pronounced differences within and between the lateral and subcortical networks. Two of the components of the lateral network, the somatosensory 2 and the insular cortex, have been shown to participate in multimodal sensory processing in both humans and rodents.^66,67^ The observed decreased FC in these regions in the zQ175DN HET mice at 12 months, is in line with previous findings in PwHD in Huntington’s disease patients in regions pertaining to the sensory-motor network.^68^ However, some clinical studies in PwHD have also shown increased FC within this network, attributing those changes to a generalized activity spread due to a loss of specialization in neuronal function.^69,70^ The lateral CPu, part of the subcortical network, also shows a reduction in FC with the insular and piriform cortex. CPu connectivity has been extensively mapped, showing compartmentalized differences in projections with both of these regions.^71,72^ Interestingly, the insular cortex has been shown to have a relevant function in switching between large-scale brain networks^73^ but also in body spatial awareness, which has been reported as altered in PwHD. ^63^ Moreover, as the piriform cortex is the key area in olfactory processing, dysfunction in olfactory discrimination is present in both PwHD^74,75^ and rodent models, specifically in the zQ175 HET model at the same age of 12 months.^76,77^

A major advantage of our longitudinal study was the ability to look at the potential disease-like modifying effects on normal aging in different large-scale brain network FC in the zQ175DN HET mice. Network changes as a consequence of normal aging have been shown in both humans and rodents.^39,78,79^ The first detailed examination of the aging effect on functional brain networks in rodents followed the DMLN, sensorimotor and subcortical networks from 3 to 13 months of age and observed an evolution in the form of an inverse U-shape.^39^ We observed the same nonlinear behavior in the WT groups, with an FC increase from 3 to 6 and later a decrease from 6 to 10 months of age. In the case of the zQ175DN HET mice, selected region pairs from each network also followed the non-linear inverted U-shape change. The regions belonging to the DMLN and the associative cortical network followed this pattern, especially in the cingulate-retrosplenial cortex FC where we observed decreased FC in the zQ175DN HET group at motor deficit onset, revealing that the disease-driven alterations impede the typical age effect of increased FC from 3 to 6 months. An overall distinct genotypic effect was observed in the interhemispheric retrosplenial-visual cortex connectivity but also between the auditory with the visual and retrosplenial cortex, reiterating the vulnerability of the DMLN and ACN regions. The normal aging effect on network FC findings followed the same nonlinear inverted U-shape as the region-based FC, especially within all networks except for the lateral network where only a significantly decreased FC from 6 to 10 months was present. In line with the region-based FC, the apparent genotype effect was present in the intra- and inter-connectivity of the DMLN and the associative cortical network. As shown from tracing studies, the strong connectivity between these two networks^80^ and between the lateral and subcortical network^71^ is also reflected in the clearly inverted trend that these network pairs followed. The lateral cortical network is the rodent homologous of the task-positive network (TPN), the anti-correlated network to the DMN^81^. We have consistently found low anti-correlated FC with DMLN and associative cortical network in the zQ175DN HET mice. Hence, this was aligned with the finding that the lateral network connectivity with DMLN and the associative cortical network showed no significant trend across time in both genotypes.

We didn’t observe any significant region or network-level FC alterations at 3 months of age in this model. Electrophysiological studies in this model have already demonstrated changes in cortical frequencies that reflect increased synaptic responses.^24^ Additionally, increased local inhibitory activity in the cortical projection neurons of the motor cortex is present as early as 2 months of age in the zQ175 HET model as a means of counteracting and preventing the cortical excitation to reach medium spiny neurons in the striatum.^23^ Similarly, cortical hyperexcitability, measured by two-photon calcium imaging, was also found before motor abnormalities appear in the R6/2 HD mouse model.^25^ Suggesting the cortex as a relevant target in Huntington’s disease is a study that shows that reduced full-length m*Htt* expression in the motor cortex of a mouse model leads to rescue in striatal activity.^82^ Cortical alterations have been reported in the homozygous form of zQ175 mice, where EEG recordings show field disruptions even before 3 months of age.^83^ Our findings imply that resting-state FC, which measures average correlation across the entire scanning duration, may not be sufficiently sensitive to pick up differences at 3 months. More advanced methods that capture temporal fluctuations in FC during the scan may be more suitable to detect more subtle changes occurring at this early age.^84,85^

Regions relevant for movement execution and control are the motor and somatosensory cortices. The zQ175DN HET mice showed more robust changes in FC in these regions in a late manifest state, at 12 months of age, albeit the initially decreased FC between retrosplenial and motor cortex at 6 months. Concerning human studies, the sensory-motor network has been implicated to follow a non-linear trajectory of pre-manifest hypo-connectivity towards a hyper-connectivity pattern in clinical stages^20^ (Fig. 7). However, many factors such as different stages of the disease, movement, medication intake, and inclusion criteria can influence the lack of reproducibility of these results in different studies.

**Figure 7.**
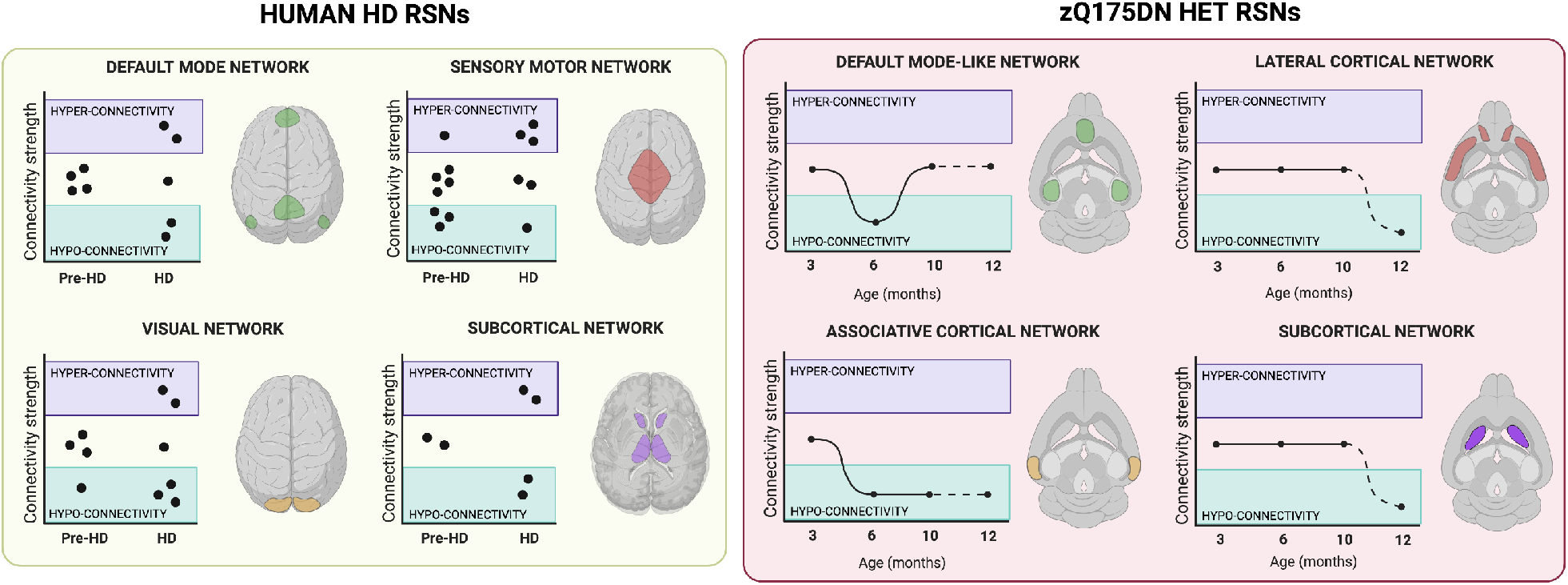
RSN alterations along the course of Huntington’s disease in humans and in the zQ175DN HET mouse model. Human rsfMRI cross-sectional studies (each point represents a study’s result) demonstrate diverse outcomes in different RSNs in both pre- and manifest stages relative to age-matched controls; Longitudinal course of RSNs evolution from pre-manifest to different manifest states in the zQ175DN HET mice compared to age-matched controls (full line links points from the longitudinal study while dashed line links the cross-sectional study point); HD = Huntington’s disease. Figure created with BioRender.com.

Although the zQ175DN HET mouse model has been shown to closely mimic several aspects of Huntington’s disease^28,29^, an important limitation is that it is still a preclinical model that, despite showing striatal and cortical volume decreases as early as 3-4 months, it does not show neuronal loss at a very early point, which is one of the hallmarks of Huntington’s disease.^86,87^ While no striking loss is identified in the zQ175 HET, there are changes in neuronal morphology.^88^ However, neurons are only one of the contributing elements of the BOLD signal in RSNs that could potentially influence the whole-brain network architecture. ^89^ Hence, an additional assessment of other components of the neurovascular unit, such as astrocytes and pericytes, could potentially elucidate the effect of mHTT on these cell populations with regards to the disease-driven network alterations.^90^

In summary, our findings demonstrate for the first time the longitudinal perturbations in large-scale brain networks that occur in a Huntington’s disease mouse model (Fig. 7). The results provide insight into how connectivity in distinct cortical and subcortical regions is divergently affected along different disease-like phenotypic states. Moreover, our data emphasize how cortical circuitries play an important role in whole-brain network changes in the zQ175 HET HD mouse model, and suggest that further research is warranted to investigate the underlying neural dynamics of these cortical disease-dependent changes. Closer investigation of the specific cellular and molecular mechanisms of different cortico-cortical and cortico-striatal circuits will provide a better understanding of the cellular network interaction in Huntington’s disease.

## References

1. Bates GP, Dorsey R, Gusella JF, et al. Huntington disease. Nat Rev Dis Primers. Apr 23 2015;1:15005. doi:10.1038/nrdp.2015.5

2. Ghosh R, Tabrizi SJ. Clinical Features of Huntington’s Disease. Adv Exp Med Biol. 2018;1049:1–28. doi:10.1007/978-3-319-71779-1_1

3. A novel gene containing a trinucleotide repeat that is expanded and unstable on Huntington’s disease chromosomes. The Huntington’s Disease Collaborative Research Group. Cell. Mar 26 1993;72(6):971–83. doi:10.1016/0092-8674(93)90585-e

4. Ross CA, Tabrizi SJ. Huntington’s disease: from molecular pathogenesis to clinical treatment. Lancet Neurol. Jan 2011;10(1):83–98. doi:10.1016/S1474-4422(10)70245-3

5. Rosas HD, Hevelone ND, Zaleta AK, Greve DN, Salat DH, Fischl B. Regional cortical thinning in preclinical Huntington disease and its relationship to cognition. Neurology. Sep 13 2005;65(5):745–7. doi:10.1212/01.wnl.0000174432.87383.87

6. Rosas HD, Salat DH, Lee SY, et al. Cerebral cortex and the clinical expression of Huntington’s disease: complexity and heterogeneity. Brain. Apr 2008;131(Pt 4):1057–68. doi:10.1093/brain/awn025

7. Waldvogel HJ, Kim EH, Thu DC, Tippett LJ, Faull RL. New Perspectives on the Neuropathology in Huntington’s Disease in the Human Brain and its Relation to Symptom Variation. J Huntingtons Dis. 2012;1(2):143–53. doi:10.3233/JHD-2012-120018

8. Estrada-Sanchez AM, Rebec GV. Role of cerebral cortex in the neuropathology of Huntington’s disease. Front Neural Circuits. 2013;7:19. doi:10.3389/fncir.2013.00019

9. Nopoulos PC, Aylward EH, Ross CA, et al. Cerebral cortex structure in prodromal Huntington disease. Neurobiol Dis. Dec 2010;40(3):544–54. doi:10.1016/j.nbd.2010.07.014

10. Unschuld PG, Joel SE, Liu X, et al. Impaired cortico-striatal functional connectivity in prodromal Huntington’s Disease. Neurosci Lett. Apr 18 2012;514(2):204–9. doi:10.1016/j.neulet.2012.02.095

11. Poudel GR, Egan GF, Churchyard A, Chua P, Stout JC, Georgiou-Karistianis N. Abnormal synchrony of resting state networks in premanifest and symptomatic Huntington disease: the IMAGE-HD study. J Psychiatry Neurosci. Mar 2014;39(2):87–96. doi:10.1503/jpn.120226

12. Tabrizi SJ, Schobel S, Gantman EC, et al. A biological classification of Huntington’s disease: the Integrated Staging System. Lancet Neurol. Jul 2022;21(7):632–644. doi:10.1016/S1474-4422(22)00120-X

13. Zeun P, Scahill RI, Tabrizi SJ, Wild EJ. Fluid and imaging biomarkers for Huntington’s disease. Mol Cell Neurosci. Jun 2019;97:67–80. doi:10.1016/j.mcn.2019.02.004

14. D JF, Stout JC, Poudel G, et al. Multimodal imaging biomarkers in premanifest and early Huntington’s disease: 30-month IMAGE-HD data. Br J Psychiatry. Jun 2016;208(6):571–8. doi:10.1192/bjp.bp.114.156588

15. Wilson H, De Micco R, Niccolini F, Politis M. Molecular Imaging Markers to Track Huntington’s Disease Pathology. Front Neurol. 2017;8:11. doi:10.3389/fneur.2017.00011

16. van den Heuvel MI, Turk E, Manning JH, et al. Hubs in the human fetal brain network. Dev Cogn Neurosci. Apr 2018;30:108–115. doi:10.1016/j.dcn.2018.02.001

17. Fransson P. Spontaneous low-frequency BOLD signal fluctuations: an fMRI investigation of the resting-state default mode of brain function hypothesis. Hum Brain Mapp. Sep 2005;26(1):15–29. doi:10.1002/hbm.20113

18. Zhang HY, Wang SJ, Liu B, et al. Resting brain connectivity: changes during the progress of Alzheimer disease. Radiology. Aug 2010;256(2):598–606. doi:10.1148/radiol.10091701

19. Luo C, Song W, Chen Q, et al. Reduced functional connectivity in early-stage drug-naive Parkinson’s disease: a resting-state fMRI study. Neurobiol Aging. Feb 2014;35(2):431–41. doi:10.1016/j.neurobiolaging.2013.08.018

20. Pini L, Jacquemot C, Cagnin A, et al. Aberrant brain network connectivity in presymptomatic and manifest Huntington’s disease: A systematic review. Hum Brain Mapp. Jan 2020;41(1):256–269. doi:10.1002/hbm.24790

21. Dumas EM, van den Bogaard SJ, Hart EP, et al. Reduced functional brain connectivity prior to and after disease onset in Huntington’s disease. Neuroimage Clin. 2013;2:377–84. doi:10.1016/j.nicl.2013.03.001

22. Wolf RC, Sambataro F, Vasic N, et al. Visual system integrity and cognition in early Huntington’s disease. Eur J Neurosci. Jul 2014;40(2):2417–26. doi:10.1111/ejn.12575

23. Indersmitten T, Tran CH, Cepeda C, Levine MS. Altered excitatory and inhibitory inputs to striatal medium-sized spiny neurons and cortical pyramidal neurons in the Q175 mouse model of Huntington’s disease. J Neurophysiol. Apr 1 2015;113(7):2953–66. doi:10.1152/jn.01056.2014

24. Donzis EJ, Estrada-Sanchez AM, Indersmitten T, et al. Cortical Network Dynamics Is Altered in Mouse Models of Huntington’s Disease. Cereb Cortex. Apr 14 2020;30(4):2372–2388. doi:10.1093/cercor/bhz245

25. Burgold J, Schulz-Trieglaff EK, Voelkl K, et al. Cortical circuit alterations precede motor impairments in Huntington’s disease mice. Sci Rep. Apr 29 2019;9(1):6634. doi:10.1038/s41598-019-43024-w

26. Sepers MD, Mackay JP, Koch E, et al. Altered cortical processing of sensory input in Huntington disease mouse models. Neurobiol Dis. Apr 20 2022;169:105740. doi:10.1016/j.nbd.2022.105740

27. Gu X, Li C, Wei W, et al. Pathological cell-cell interactions elicited by a neuropathogenic form of mutant Huntingtin contribute to cortical pathogenesis in HD mice. Neuron. May 5 2005;46(3):433–44. doi:10.1016/j.neuron.2005.03.025

28. Menalled LB, Kudwa AE, Miller S, et al. Comprehensive behavioral and molecular characterization of a new knock-in mouse model of Huntington’s disease: zQ175. PLoS One. 2012;7(12):e49838. doi:10.1371/journal.pone.0049838

29. Heikkinen T, Bragge T, Bhattarai N, et al. Rapid and robust patterns of spontaneous locomotor deficits in mouse models of Huntington’s disease. PLoS One. 2020;15(12):e0243052. doi:10.1371/journal.pone.0243052

30. Carty N, Berson N, Tillack K, et al. Characterization of HTT inclusion size, location, and timing in the zQ175 mouse model of Huntington’s disease: an in vivo high-content imaging study. PLoS One. 2015;10(4):e0123527. doi:10.1371/journal.pone.0123527

31. Southwell AL, Smith-Dijak A, Kay C, et al. An enhanced Q175 knock-in mouse model of Huntington disease with higher mutant huntingtin levels and accelerated disease phenotypes. Hum Mol Genet. Sep 1 2016;25(17):3654–3675. doi:10.1093/hmg/ddw212

32. Fox JG BS, Davisson MT, Newcomer CE, Quimby FW, Smith AL, eds. The Mouse in Biomedical Research, 2nd Edition. vol 3. Elsevier 2007:637–72.

33. Jonckers E, Delgado y Palacios R, Shah D, Guglielmetti C, Verhoye M, Van der Linden A. Different anesthesia regimes modulate the functional connectivity outcome in mice. Magn Reson Med. Oct 2014;72(4):1103–12. doi:10.1002/mrm.24990

34. Grandjean J, Schroeter A, Batata I, Rudin M. Optimization of anesthesia protocol for resting-state fMRI in mice based on differential effects of anesthetics on functional connectivity patterns. Neuroimage. Nov 15 2014;102 Pt 2:838–47. doi:10.1016/j.neuroimage.2014.08.043

35. Praet J, Manyakov NV, Muchene L, et al. Diffusion kurtosis imaging allows the early detection and longitudinal follow-up of amyloid-beta-induced pathology. Alzheimers Res Ther. Jan 9 2018;10(1):1. doi:10.1186/s13195-017-0329-8

36. Liska A, Galbusera A, Schwarz AJ, Gozzi A. Functional connectivity hubs of the mouse brain. Neuroimage. Jul 15 2015;115:281–91. doi:10.1016/j.neuroimage.2015.04.033

37. Bukhari Q, Schroeter A, Cole DM, Rudin M. Resting State fMRI in Mice Reveals Anesthesia Specific Signatures of Brain Functional Networks and Their Interactions. Front Neural Circuits. 2017;11:5. doi:10.3389/fncir.2017.00005

38. Janke AL, Ullmann JF. Robust methods to create ex vivo minimum deformation atlases for brain mapping. Methods. Feb 2015;73:18–26. doi:10.1016/j.ymeth.2015.01.005

39. Egimendia A, Minassian A, Diedenhofen M, Wiedermann D, Ramos-Cabrer P, Hoehn M. Aging Reduces the Functional Brain Networks Strength-a Resting State fMRI Study of Healthy Mouse Brain. Front Aging Neurosci. 2019; 11:277. doi:10.3389/fnagi.2019.00277

40. Gozzi A, Schwarz AJ. Large-scale functional connectivity networks in the rodent brain. Neuroimage. Feb 15 2016;127:496–509. doi:10.1016/j.neuroimage.2015.12.017

41. Sforazzini F, Schwarz AJ, Galbusera A, Bifone A, Gozzi A. Distributed BOLD and CBV-weighted resting-state networks in the mouse brain. Neuroimage. Feb 15 2014;87:403–15. doi:10.1016/j.neuroimage.2013.09.050

42. Raichle ME. The brain’s default mode network. Annu Rev Neurosci. Jul 8 2015;38:433–47. doi:10.1146/annurev-neuro-071013-014030

43. Buckner RL, Andrews-Hanna JR, Schacter DL. The brain’s default network: anatomy, function, and relevance to disease. Ann N Y Acad Sci. Mar 2008; 1124:1–38. doi:10.1196/annals.1440.011

44. Jones DT, Knopman DS, Gunter JL, et al. Cascading network failure across the Alzheimer’s disease spectrum. Brain. Feb 2016;139(Pt 2):547–62. doi:10.1093/brain/awv338

45. Koch W, Teipel S, Mueller S, et al. Diagnostic power of default mode network resting state fMRI in the detection of Alzheimer’s disease. Neurobiol Aging. Mar 2012;33(3):466–78. doi:10.1016/j.neurobiolaging.2010.04.013

46. van Eimeren T, Monchi O, Ballanger B, Strafella AP. Dysfunction of the default mode network in Parkinson disease: a functional magnetic resonance imaging study. Arch Neurol. Jul 2009;66(7):877–83. doi:10.1001/archneurol.2009.97

47. Laubach M, Amarante LM, Swanson K, White SR. What, If Anything, Is Rodent Prefrontal Cortex? eNeuro. Sep-Oct 2018;5(5)doi:10.1523/ENEURO.0315-18.2018

48. Quarantelli M, Salvatore E, Giorgio SM, et al. Default-mode network changes in Huntington’s disease: an integrated MRI study of functional connectivity and morphometry. PLoS One. 2013;8(8):e72159. doi:10.1371/journal.pone.0072159

49. Hobbs NZ, Pedrick AV, Say MJ, et al. The structural involvement of the cingulate cortex in premanifest and early Huntington’s disease. Mov Disord. Aug 1 2011;26(9):1684–90. doi:10.1002/mds.23747

50. Yamawaki N, Radulovic J, Shepherd GM. A Corticocortical Circuit Directly Links Retrosplenial Cortex to M2 in the Mouse. J Neurosci. Sep 7 2016;36(36):9365–74. doi:10.1523/JNEUROSCI.1099-16.2016

51. Opalka AN, Wang DV. Hippocampal efferents to retrosplenial cortex and lateral septum are required for memory acquisition. Learn Mem. Aug 2020;27(8):310–318. doi:10.1101/lm.051797.120

52. Farrar AM, Murphy CA, Paterson NE, et al. Cognitive deficits in transgenic and knock-in HTT mice parallel those in Huntington’s disease. J Huntingtons Dis. 2014;3(2):145–58. doi:10.3233/JHD-130061

53. Lione LA, Carter RJ, Hunt MJ, Bates GP, Morton AJ, Dunnett SB. Selective discrimination learning impairments in mice expressing the human Huntington’s disease mutation. J Neurosci. Dec 1 1999;19(23):10428–37.

54. Paulsen JS, Conybeare RA. Cognitive changes in Huntington’s disease. Adv Neurol. 2005;96:209–25.

55. Beglinger LJ, Nopoulos PC, Jorge RE, et al. White matter volume and cognitive dysfunction in early Huntington’s disease. Cogn Behav Neurol. Jun 2005;18(2):102–7. doi:10.1097/01.wnn.0000152205.79033.73

56. Montoya A, Pelletier M, Menear M, Duplessis E, Richer F, Lepage M. Episodic memory impairment in Huntington’s disease: a meta-analysis. Neuropsychologia. 2006;44(10): 1984–94. doi:10.1016/j.neuropsychologia.2006.01.015

57. Lange KW, Sahakian BJ, Quinn NP, Marsden CD, Robbins TW. Comparison of executive and visuospatial memory function in Huntington’s disease and dementia of Alzheimer type matched for degree of dementia. J Neurol Neurosurg Psychiatry. May 1995;58(5):598–606. doi:10.1136/jnnp.58.5.598

58. Lawrence AD, Sahakian BJ, Hodges JR, Rosser AE, Lange KW, Robbins TW. Executive and mnemonic functions in early Huntington’s disease. Brain. Oct 1996;119 (Pt 5):1633–45. doi:10.1093/brain/119.5.1633

59. Albert MS, Butters N, Brandt J. Development of remote memory loss in patients with Huntington’s disease. J Clin Neuropsychol. May 1981;3(1):1–12. doi:10.1080/01688638108403109

60. Begeti F, Schwab LC, Mason SL, Barker RA. Hippocampal dysfunction defines disease onset in Huntington’s disease. J Neurol Neurosurg Psychiatry. Sep 2016;87(9):975–81. doi:10.1136/jnnp-2015-312413

61. Profant O, Roth J, Bures Z, et al. Auditory dysfunction in patients with Huntington’s disease. Clin Neurophysiol. Oct 2017;128(10):1946–1953. doi:10.1016/j.clinph.2017.07.403

62. Saft C, Schuttke A, Beste C, Andrich J, Heindel W, Pfleiderer B. fMRI reveals altered auditory processing in manifest and premanifest Huntington’s disease. Neuropsychologia. Apr 2008;46(5):1279–89. doi:10.1016/j.neuropsychologia.2007.12.002

63. Labuschagne I, Cassidy AM, Scahill RI, et al. Visuospatial Processing Deficits Linked to Posterior Brain Regions in Premanifest and Early Stage Huntington’s Disease. J Int Neuropsychol Soc. Jul 2016;22(6):595–608. doi:10.1017/S1355617716000321

64. Todd TP, Mehlman ML, Keene CS, DeAngeli NE, Bucci DJ. Retrosplenial cortex is required for the retrieval of remote memory for auditory cues. Learn Mem. Jun 2016;23(6):278–88. doi:10.1101/lm.041822.116

65. Hackett TA. Information flow in the auditory cortical network. Hear Res. Jan 2011;271(1-2):133–46. doi:10.1016/j.heares.2010.01.011

66. Downar J, Crawley AP, Mikulis DJ, Davis KD. A multimodal cortical network for the detection of changes in the sensory environment. Nat Neurosci. Mar 2000;3(3):277–83. doi:10.1038/72991

67. Rodgers KM, Benison AM, Klein A, Barth DS. Auditory, somatosensory, and multisensory insular cortex in the rat. Cereb Cortex. Dec 2008;18(12):2941–51. doi:10.1093/cercor/bhn054

68. Muller HP, Gorges M, Gron G, et al. Motor network structure and function are associated with motor performance in Huntington’s disease. J Neurol. Mar 2016;263(3):539–49. doi:10.1007/s00415-015-8014-y

69. Werner CJ, Dogan I, Sass C, et al. Altered resting-state connectivity in Huntington’s disease. Hum Brain Mapp. Jun 2014;35(6):2582–93. doi:10.1002/hbm.22351

70. Sanchez-Castaneda C, de Pasquale F, Caravasso CF, et al. Resting-state connectivity and modulated somatomotor and default-mode networks in Huntington disease. CNS Neurosci Ther. Jun 2017;23(6):488–497. doi:10.1111/cns.12701

71. Hintiryan H, Foster NN, Bowman I, et al. The mouse cortico-striatal projectome. Nat Neurosci. Aug 2016;19(8):1100–14. doi:10.1038/nn.4332

72. Gehrlach DA, Weiand C, Gaitanos TN, et al. A whole-brain connectivity map of mouse insular cortex. Elife. Sep 17 2020;9 doi:10.7554/eLife.55585

73. Menon V, Uddin LQ. Saliency, switching, attention and control: a network model of insula function. Brain Struct Funct. Jun 2010;214(5-6):655–67. doi:10.1007/s00429-010-0262-0

74. Nordin S, Paulsen JS, Murphy C. Sensory- and memory-mediated olfactory dysfunction in Huntington’s disease. J Int Neuropsychol Soc. May 1995;1(3):281–90. doi:10.1017/s1355617700000278

75. Hamilton JM, Murphy C, Paulsen JS. Odor detection, learning, and memory in Huntington’s disease. J Int Neuropsychol Soc. Nov 1999;5(7):609–15. doi:10.1017/s1355617799577035

76. Lazic SE, Goodman AO, Grote HE, et al. Olfactory abnormalities in Huntington’s disease: decreased plasticity in the primary olfactory cortex of R6/1 transgenic mice and reduced olfactory discrimination in patients. Brain Res. Jun 2 2007;1151:219–26. doi:10.1016/j.brainres.2007.03.018

77. Ferris CF, Kulkarni P, Toddes S, Yee J, Kenkel W, Nedelman M. Studies on the Q175 Knock-in Model of Huntington’s Disease Using Functional Imaging in Awake Mice: Evidence of Olfactory Dysfunction. Front Neurol. 2014;5:94. doi:10.3389/fneur.2014.00094

78. Betzel RF, Byrge L, He Y, Goni J, Zuo XN, Sporns O. Changes in structural and functional connectivity among resting-state networks across the human lifespan. Neuroimage. Nov 15 2014;102 Pt 2:345–57. doi:10.1016/j.neuroimage.2014.07.067

79. Bo J, Lee CM, Kwak Y, et al. Lifespan differences in cortico-striatal resting state connectivity. Brain Connect. Apr 2014;4(3):166–80. doi:10.1089/brain.2013.0155

80. Whitesell JD, Liska A, Coletta L, et al. Regional, Layer, and Cell-Type-Specific Connectivity of the Mouse Default Mode Network. Neuron. Feb 3 2021;109(3):545–559 e8. doi:10.1016/j.neuron.2020.11.011

81. Fox MD, Snyder AZ, Vincent JL, Corbetta M, Van Essen DC, Raichle ME. The human brain is intrinsically organized into dynamic, anticorrelated functional networks. Proc Natl Acad Sci U S A. Jul 5 2005;102(27):9673–8. doi:10.1073/pnas.0504136102

82. Estrada-Sanchez AM, Burroughs CL, Cavaliere S, et al. Cortical efferents lacking mutant huntingtin improve striatal neuronal activity and behavior in a conditional mouse model of Huntington’s disease. J Neurosci. Mar 11 2015;35(10):4440–51. doi:10.1523/JNEUROSCI.2812-14.2015

83. Fisher SP, Schwartz MD, Wurts-Black S, et al. Quantitative Electroencephalographic Analysis Provides an Early-Stage Indicator of Disease Onset and Progression in the zQ175 Knock-In Mouse Model of Huntington’s Disease. Sleep. Feb 1 2016;39(2):379–91. doi:10.5665/sleep.5448

84. Belloy ME, Shah D, Abbas A, et al. Quasi-Periodic Patterns of Neural Activity improve Classification of Alzheimer’s Disease in Mice. Sci Rep. Jul 3 2018;8(1):10024. doi:10.1038/s41598-018-28237-9

85. Adhikari MH, Belloy ME, Van der Linden A, Keliris GA, Verhoye M. Resting-State Co-activation Patterns as Promising Candidates for Prediction of Alzheimer’s Disease in Aged Mice. Front Neural Circuits. 2020;14:612529. doi:10.3389/fncir.2020.612529

86. Cowan CM, Raymond LA. Selective neuronal degeneration in Huntington’s disease. Curr Top Dev Biol. 2006;75:25–71. doi:10.1016/S0070-2153(06)75002-5

87. Heikkinen T, Lehtimaki K, Vartiainen N, et al. Characterization of neurophysiological and behavioral changes, MRI brain volumetry and 1H MRS in zQ175 knock-in mouse model of Huntington’s disease. PLoS One. 2012;7(12):e50717. doi:10.1371/journal.pone.0050717

88. Goodliffe JW, Song H, Rubakovic A, et al. Differential changes to D1 and D2 medium spiny neurons in the 12-month-old Q175+/-mouse model of Huntington’s Disease. PLoS One. 2018;13(8):e0200626. doi:10.1371/journal.pone.0200626

89. Drew PJ. Vascular and neural basis of the BOLD signal. Curr Opin Neurobiol. Oct 2019;58:61–69. doi:10.1016/j.conb.2019.06.004

90. Attwell D, Buchan AM, Charpak S, Lauritzen M, Macvicar BA, Newman EA. Glial and neuronal control of brain blood flow. Nature. Nov 11 2010;468(7321):232–43. doi:10.1038/nature09613

