## Supplementary Information for "Resting-State fMRI reveals Longitudinal Alterations in Brain Network Connectivity in a Mouse Model of Huntington’s Disease"

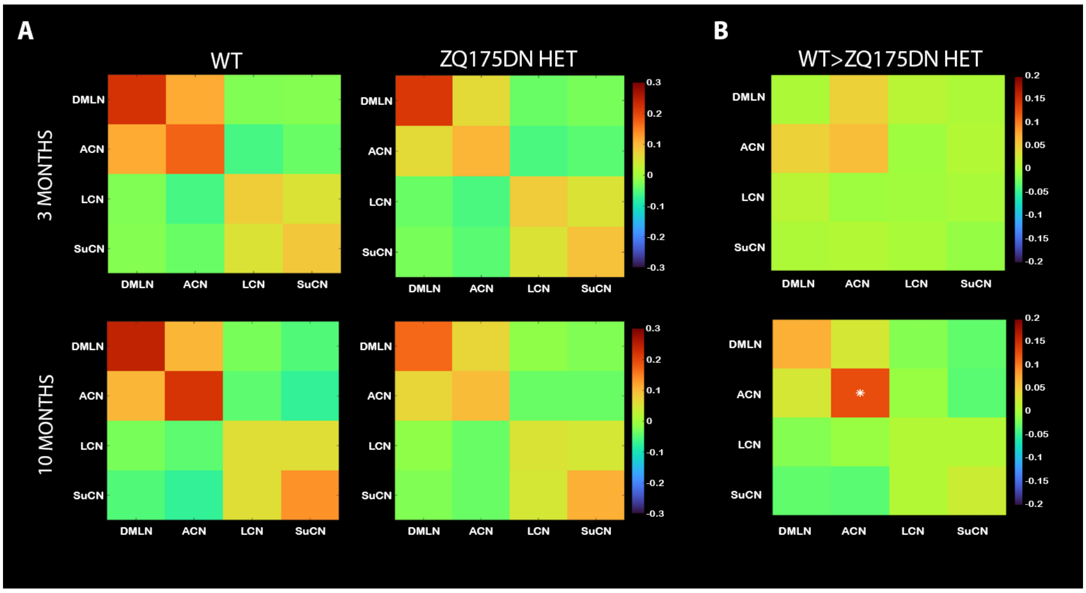


**<u>Figure S1</u>. Network FC at 3 and 10 months of age in zQ175DN HET and WT mice. (A)** Mean z-transformed correlation (mirrored) matrices of each group at 3 (top row) and 10 months of age (bottom row) – within network FC is shown along the diagonal of the matrix; red/orange colors represent positively correlated connectivity, green color indicates very low to no connectivity between regions and dark/light blue colors represent negatively correlated connectivity. **(B)** Between-group differences for 3 and 10 months of age. Red/orange represents lower positive or higher negative FC between a pair of regions in zQ175DN HET compared to WT while dark/light blue represents higher positive FC or lower negative FC in zQ175DN HET compared to WT. Asterisk indicates significant group differences in FC based on a two-sample T-test (p≤0.05, FDR corrected) only performed on connections that demonstrated a significant FC in at least one group based on a one-sample T-test.

| | | | | | 3-6M | | | | 3-10M | | | | 6-10M | | | |
|---|---|---|---|---|---|---|---|---|---|---|---|---|---|---|---|---|
| FC Pair | Genotype | Age | Genotype* Time | group | p value | Diff | lower 95% | upper 95% | p value | Diff | lower 95% | upper 95% | p value | Diff | lower 95% | upper 95% |
| RspCtx (L-R) | **0,0001** | **0,0001** | 0,1785 | | **0,0176** | -0,1532 | -0,2841 | -0,0223 | 0,1626 | 0,1011 | -0,0298 | 0,2320 | **3,70E-05** | 0,2542 | 0,1233 | 0,3851 |
| S2Ctx (L-R) | **0,0171** | **0,0057** | 0,7250 | | 0,6998 | -0,0475 | -0,1879 | 0,0929 | 0,0508 | 0,1400 | -0,0004 | 0,2804 | **0,0056** | 0,1875 | 0,0471 | 0,3280 |
| VCtx (L-R) | **5,0E-06** | **0,0001** | 0,6625 | | **0,0002** | -0,1833 | -0,2864 | -0,0802 | 0,7272 | -0,033 | -0,136 | 0,0701 | **0,0022** | 0,1504 | 0,0473 | 0,2534 |
| ICPu (L-R) | 0,7144 | **0,004** | 0,9990 | | **0,0037** | -0,1735 | -0,2954 | -0,0516 | 0,6150 | -0,0472 | -0,1691 | 0,0747 | **0,0409** | 0,1263 | 0,0044 | 0,2482 |
| Cg-RspCtx (L) | **0,0076** | **1,90E-06** | **0,0461**[a] | **WT** | **3,00E-05** | -0,3322 | -0,5166 | -0,1477 | 1 | -0,0074 | -0,1919 | 0,1770 | **4,60E-05** | 0,3247 | 0,1403 | 0,5092 |
| | | | | **HET** | 0,3171 | -0,1258 | -0,3052 | 0,0536 | 0,7761 | 0,0797 | -0,0997 | 0,2591 | **0,016** | 0,2055 | 0,0261 | 0,3849 |
| Au-VCtx (L) | **0,0001** | 0,1218 | 0,1947 | | / | / | / | / | / | / | / | / | / | / | / | / |
| Au-RspCtx (L) | **0,0018** | **0,0011** | 0,1289 | | **0,0115** | -0,0904 | -0,1637 | -0,0171 | 0,7879 | 0,0203 | -0,0530 | 0,0935 | **0,0015** | 0,1106 | 0,0373 | 0,1839 |
| S2-MCtx (L) | 0,5309 | 0,1285 | 0,9406 | | / | / | / | | / | / | / | / | / | / | / | / |

<u>Table 2</u>. LMM statistics based on ROI FC

| | | | | 3-6M | | | | 3-10M | | | | 6-10M | | | |
|---|---|---|---|---|---|---|---|---|---|---|---|---|---|---|---|
| **FC Network** | **Genotype** | **Age** | **Genotype *Time** | **p value** | **Diff** | **lower 95%** | **upper 95%** | **p value** | **Diff** | **lower 95%** | **upper 95%** | **p value** | **Diff** | **lower 95%** | **upper 95%** |
| wDMLN | **0,0038** | **4,40E-08** | 0,0805 | **2,30E-06** | -0,1596 | -0,2311 | -0,0880 | 0,8868 | 0,0140 | -0,0575 | 0,0856 | **3,20E-07** | 0,1736 | 0,1020 | 0,2451 |
| wACN | **2,50E-06** | **0,0009** | 0,3886 | **0,0009** | -0,0939 | -0,1537 | -0,0341 | 0,6099 | -0,0239 | -0,0837 | 0,0359 | **0,0174** | 0,0701 | 0,0103 | 0,1299 |
| wLCN | 0,5339 | **0,0012** | 0,8849 | 0,1552 | -0,0257 | -0,0586 | 0,0072 | 0,1282 | 0,0270 | -0,0058 | 0,0599 | **0,0007** | 0,0527 | 0,0199 | 0,0856 |
| wSuCN | 0,7025 | **2,30E-05** | 0,7882 | **1,60E-05** | -0,08621 | -0,1258 | -0,0466 | 0,1528 | -0,0308 | -0,0704 | 0,0088 | **0,0044** | 0,0554 | 0,0158 | 0,0950 |
| DMLN-ACN | **0,0003** | **2,70E-07** | 0,2201 | **4,70E-06** | -0,0955 | -0,1397 | -0,0512 | 0,9953 | 0,0017 | -0,0425 | 0,0459 | **3,20E-06** | 0,0972 | 0,0530 | 0,1414 |
| LCN-SuCN | 0,3978 | **0,0004** | 0,9538 | **0,0030** | -0,0359 | -0,0606 | -0,0112 | 0,8865 | 0,0047 | -0,0200 | 0,0294 | **0,0008** | 0,0406 | 0,0159 | 0,0653 |
| DMLN-LCN | 0,6805 | 0,0927 | 0,4189 | / | / | / | / | / | / | / | / | / | / | / | / |
| LCN-ACN | 0,8134 | 0,1897 | 0,6294 | / | / | / | / | / | / | / | / | / | / | / | / |

Table 3. LMM statistics based on Network FC
